# Gjd2b-mediated gap junctions promote glutamatergic synapse formation and dendritic elaboration in Purkinje neurons

**DOI:** 10.1101/2020.01.31.928242

**Authors:** Sahana Sitaraman, Gnaneshwar Yadav, Shaista Jabeen, Vandana Agarwal, Vatsala Thirumalai

**Author notes:** These authors contributed equally (Sahana Sitaraman, Gnaneshwar Yadav).

## Abstract

Gap junctions between neurons serve as electrical synapses, in addition to conducting metabolites and signaling molecules. These functions of gap junctions have led to the idea that during development, gap junctions could prefigure chemical synapses. We present evidence for this idea at a central, glutamatergic synapse and provide some mechanistic insights. Here, we show that reduction or loss of Gjd2b-containing gap junctions led to a decrease in glutamatergic synapse density in cerebellar Purkinje neurons (PNs) in larval zebrafish. Gjd2b^-/-^ larvae exhibited faster mEPSCs and a consistent decrease in dendritic arbor size. These PNs also showed decreased branch elongations but normal rate of branch retractions. Further, the dendritic growth deficits in gjd2b^-/-^ mutants were rescued by expressing full length Gjd2b in single PNs. This suggests that Gjd2b may form heterotypic channels with other connexins in gjd2b^-/-^ larvae, though it is not clear if PNs in wild type animals make homotypic or heterotypic gap junction channels. Dendritic growth deficits were not rescued by expressing a deletion mutant of Gjd2b unable to form functional channels. Finally, the expression levels of five isoforms of *camkii* were increased in gjd2b^-/-^ larvae and inhibition of CaMKII restored dendritic arbor lengths of mutant larvae to wild type levels. These results suggest a link between signaling via Gjd2b-containing gap junctions, CaMKII function and dendritic growth. In sum, our results demonstrate that Gjd2b-mediated gap junctions are key regulators of glutamatergic synapse formation and dendritic elaboration in PNs.

## Introduction

The formation of synapses is an elaborate multi-step process involving several classes of signaling molecules and electrical activity [1, 2]. Several studies support the view that connections that favor correlated activity between presynaptic and post-synaptic neurons are strengthened while those that are poorly correlated are weakened and eliminated [3, 4]. Correlated activity promotes dendritic elaboration [5, 6] and the subsequent formation of synaptic sites on those newly formed arbors [7]. However, we understand little about the molecular mechanisms linking correlated activity to dendritic arbor elaboration and synaptogenesis [8–11].

Electrical synapses, formed via gap junctions between connected neuronal pairs, allow the passage of ions, metabolites and second messengers and are ideally suited for enhancing correlations in activity between connected neurons [12, 13]. Indeed, several lines of evidence suggest that gap junctions play critical roles in circuit assembly. First, neurons show increased gap junctional connectivity early on in development at stages that precede chemical synapse formation [14–16]. Second, knocking out or knocking down gap junction proteins at these stages results in decreased chemical synapse connectivity at later stages [17, 18]. Gap junctions could mediate chemical synaptogenesis by increasing correlations in activity, by transmitting synaptogenic signaling molecules, or by providing enhanced mechanical stability at junctional sites between connected pairs. It is not clear which of these functions of gap junctions are critical for synaptogenesis. In addition, while the role of excitatory chemical synapses in sculpting neuronal arbors has been investigated in several circuits [19–21], little is known regarding such a role for electrical synapses. We set out to investigate whether gap junctions regulate structural and functional synaptic development of cerebellar Purkinje neurons and if yes, what mechanisms may be involved using larval zebrafish as our model system.

The cerebellum is critical for maintaining balance and for co-ordination of movements. It is one of the most primitive organs of the vertebrate central nervous system, has a layered structure and its circuitry is conserved from fish to mammals [22]. Purkinje neurons (PNs) are principal output neurons of the cerebellar cortex and receive excitatory and inhibitory synaptic inputs on their elaborate dendritic arbors. PNs receive thousands of glutamatergic inputs from parallel fiber axons of granule cells and relatively fewer inhibitory inputs from molecular layer interneurons. They also receive strong excitatory inputs from inferior olivary climbing fiber axons on their proximal dendrites. In zebrafish, PNs are specified by 2.5 days post fertilization (dpf), begin elaborating dendritic arbors soon after and a distinct molecular layer consisting of PN dendritic arbors becomes visible by 5 dpf [23, 24]. In addition, excitatory and inhibitory synaptic currents can be recorded in PNs of 4 dpf zebrafish larvae, evidencing a nascent functional circuit at this stage [25].

The gap junction protein Gjd2b, also referred to as Connexin 35b or 35.1 (Cx35/Cx35.1) is the teleostean homolog of the mammalian Cx36, which is the predominant neural gap junction protein. PNs begin to express Gjd2b by about 4 dpf and the level of expression steadily increases at least until 15 dpf [16]. We asked whether Gjd2b regulates structural and functional synaptic development of PNs. Using knock-down and knock-out approaches, we show here that Gjd2b is indeed required for the formation of glutamatergic synapses on PNs; that Gjd2b is also required for normal dendritic arbor growth via a CaMKII dependent process; that post-synaptic Gjd2b in PNs is sufficient to direct these processes, but that functional Gjd2b channels are necessary for such rescue. These results together suggest that signaling via Gjd2b-containing electrical synapses leads to long-lasting changes in CaMKII levels culminating in dendritic growth and subsequently, the formation of new synapses on these nascent branches.

## Results

The gap junction protein Gjd2b is expressed at high levels in the cerebellum of larval zebrafish beginning at 4 dpf, a stage at which cerebellar neurons have been specified but chemical synaptic connections are still forming [23, 16] (Figure S1A). Further we confirmed that Gjd2b puncta localized to PN cell membrane in their cell bodies and dendrites (Figure S1B). To test whether Gjd2b mediates chemical synaptogenesis in PNs, we knocked it down with a splice-blocking morpholino antisense oligonucleotide targeted to the splice junction between exon1 and intron1 of gjd2b (Figure S2A, top). Injection of this splice blocking morpholino (SBMO) into 1- 4 cell stage embryos interferes with normal splicing of the gjd2b gene product, resulting in the inclusion of a 40 base intronic segment in the mature mRNA (Figure S2C, left). The mis-spliced gene product was detected in 2 dpf and 5 dpf larvae (Figure S2B) and the sequence reveals a premature stop codon resulting in a putative truncated protein of 24 amino acid residues in the N-terminus (Figure S2C, right). Gjd2b immunoreactivity in the PN and molecular layers of the cerebellum is reduced in SBMO larvae compared to uninjected larvae (Figure S2D) confirming effective knock-down of Gjd2b levels. Injection of a morpholino with a 5-base mismatch (CTRL) did not alter Gjd2b protein levels (Figure S2E).

We recorded AMPAR-mediated miniature excitatory post-synaptic currents in PNs of uninjected and SBMO larvae using previously established methods [25]. mEPSCs in morphants occurred less frequently than in uninjected larvae (Figure 1A), reflected as increased inter-event intervals in the morphants (Figure 1C). mEPSCs in morphants also showed a small increase in the peak amplitude (Figure 1D) and faster decay time constants (Figure 1B and 1F).

**Figure 1:**
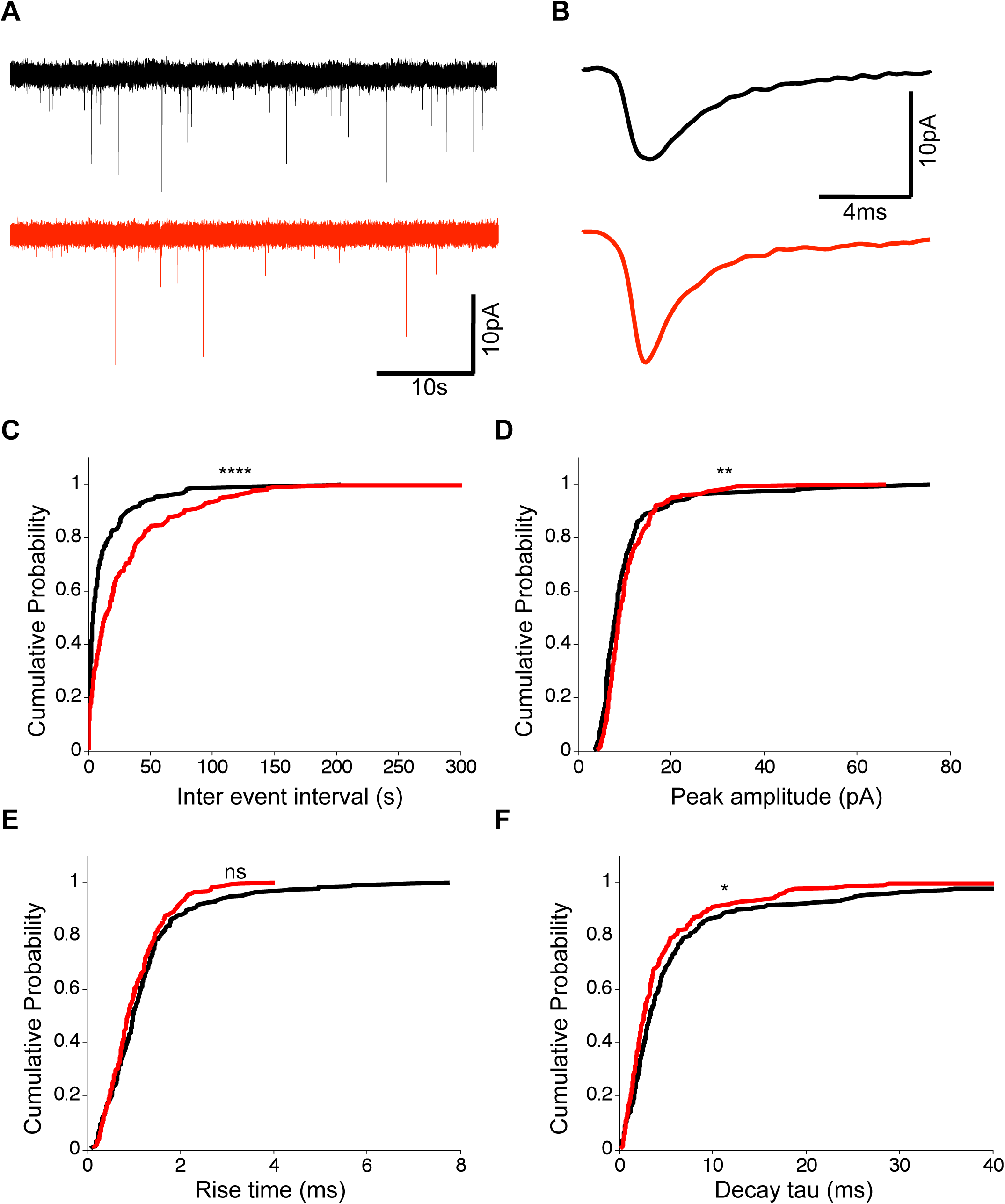
Knocking down Gjd2b reduces glutamatergic mEPSC frequency in PNs by potentially decreasing synaptic number. A: Raw traces of mEPSC recordings from PNs in control (black) and SBMO injected (red) larvae. B: Average mEPSC waveforms from control and SBMO injected larvae. C-F: Cumulative probability distributions of mEPSC inter-event intervals (C), peak amplitudes (D), 10- 90% rise times (E) and decay tau (F) in control and SBMO injected larvae. N = 7 cells from 7 larvae for control and 12 cells from 12 larvae for the SBMO group. **P<0.01; ***P<0.001; ****P<0.0001; Mann Whitney U test. See also Figures S1 and S2.

The increase in inter-event intervals of mEPSCs suggests that knocking down Gjd2b leads to a decrease in the density of synapses impinging on PNs. However, this change may also be due to a decrease in presynaptic probability of transmitter release or an increase in the number of NMDAR-only ‘silent’ synapses [26, 27]. To test if these possibilities are likely, we recorded synaptic currents evoked in wild type PNs after stimulation of climbing fibers (CFs; Figure 2A). As is the case in developing mammalian PNs [28], we recorded no NMDAR-mediated component of the evoked EPSC in PNs (Figure 2B), suggesting that the decrease in mEPSC frequency after Gjd2b knock-down is likely not due to silent synapses. Secondly, we measured paired pulse ratios (PPR) as an indicator of changes in presynaptic vesicle release probability. The CF-PN synapse shows paired-pulse depression when pulses are placed 35 ms apart (Figure 2C). We tested a range of inter-stimulus intervals (ISIs) from 30 ms till 550 ms and found that the PPR varied as a function of the ISI but did not vary significantly between the uninjected and SBMO groups (Figure 2D). Taken together, these results suggest that knocking down Gjd2b leads to a decrease in the number of glutamatergic synapses impinging on PNs.

**Figure 2:**
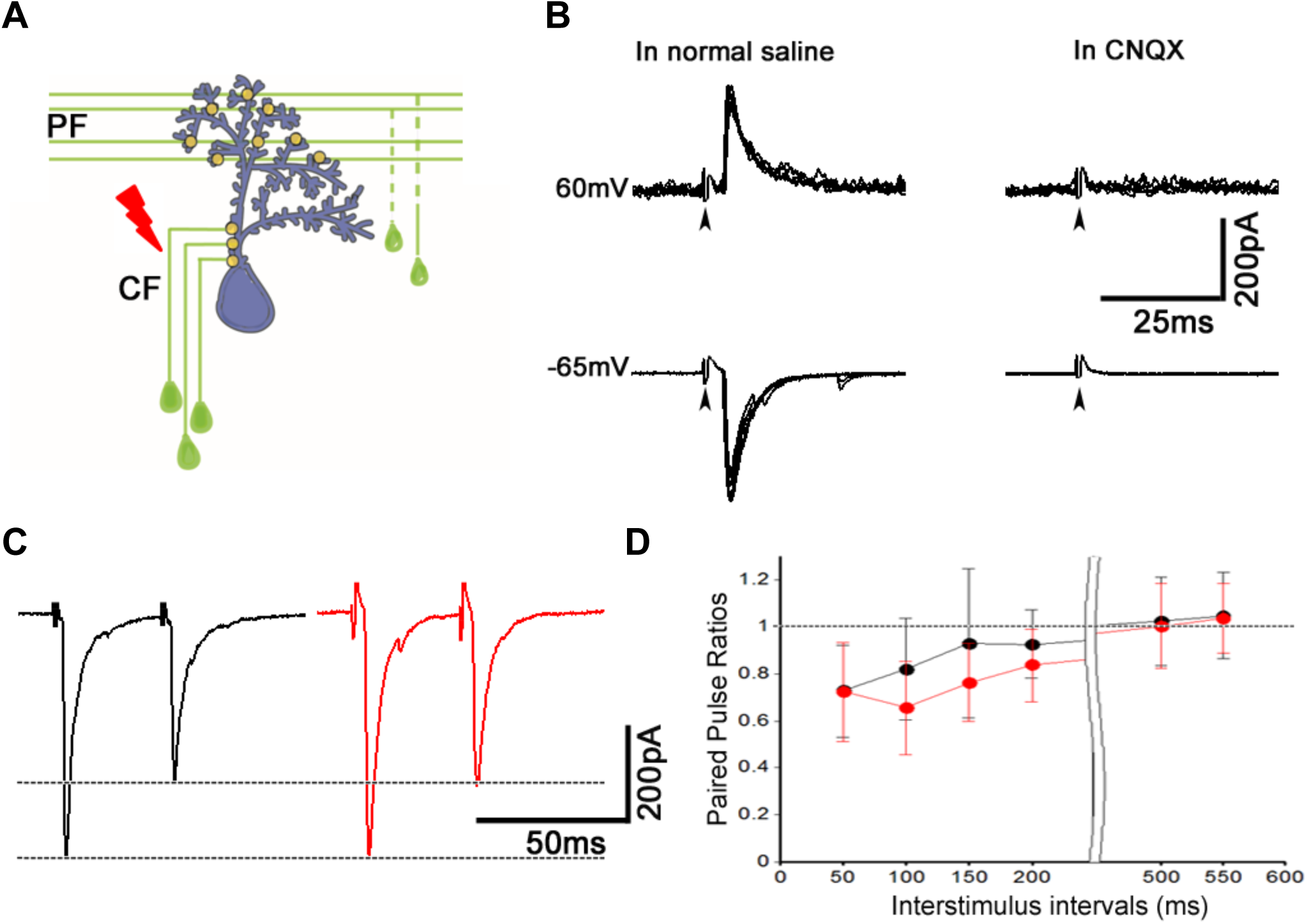
Reduction in mEPSC frequency in Gjd2b-KD animals is not due to silent synapses or change in paired pulse ratio. A: Schematic of experimental setup for stimulating climbing fibers (CF) while recording EPSCs in PNs (blue). B: EPSCs recorded at a hyperpolarized holding potential (bottom row traces) and at a depolarized holding potential (top traces) in normal saline (left side traces) and in saline containing the AMPAR blocker CNQX (right side traces). No EPSCs were detected in the presence of CNQX at −65 or +60mV. N = 6 cells, mixed ages. C: Paired pulse depression of EPSCs in PNs of control (black) and SBMO injected (red) larvae. D: Paired pulse ratios were not significantly different between control and SBMO injected larvae at any of the interstimulus intervals tested (Mean±SEM; N= 5 cells from 5 larvae each in control and SBMO groups; Two-way repeated measures ANOVA, P=0.081 for groups (control, SBMO) and P<0.001 for ISI). See also Figures S1 and S2.

As knockdown with morpholinos has been shown to sometimes result in off-target and non-specific effects [29, 30], we generated gjd2b^-/-^ fish using TALENs [31] targeted against exon1 of gjd2b (Figure 3A). We isolated several alleles with indels in the target region and chose one allele, gjd2b^ncb215^ (hereinafter referred to as gjd2b^-/-^) for further analysis. In this allele, insertion of a single G after the translation start site abolished an *XhoI* restriction enzyme recognition site within the target region (Figure 3B), and resulted in failure of *XhoI* restriction digestion in homozygotes and partial digestion in heterozygotes (Figure 3C). This single nucleotide insertion caused a frame shift and insertion of a premature stop codon, with a predicted truncated protein of 55 amino acid residues (Figure 3D). Homozygotes with this mutation showed reduced Gjd2b-like immunoreactivity in their cerebellum (Figure 3E).

**Figure 3:**
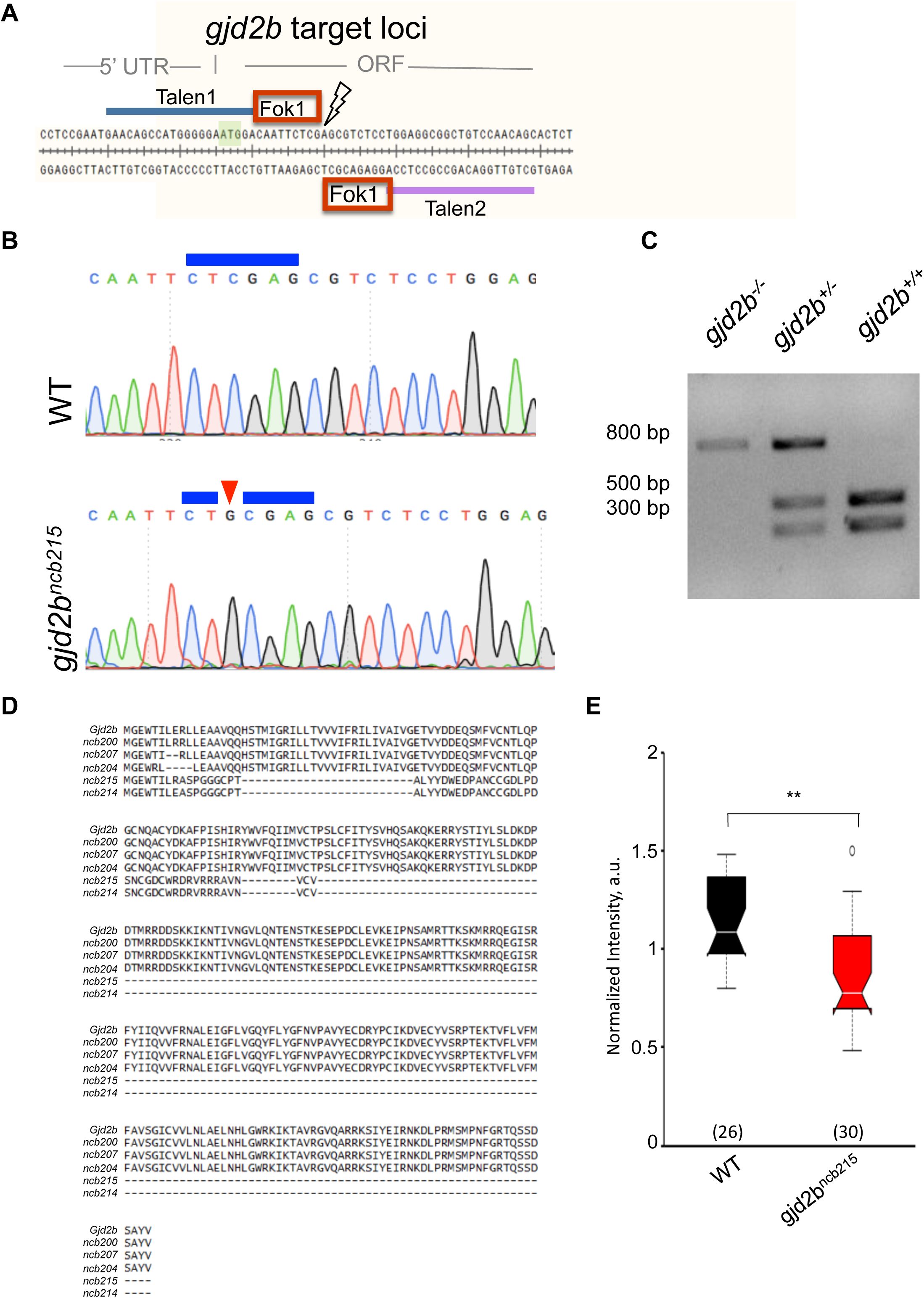
Generation of gjd2b mutant zebrafish. A: Genomic region around the start codon (green box) of gjd2b gene was selected for TALEN design, where TALEN-1 recognition sequence (red line) spans the 5’UTR, the start codon sequence and a few nucleotides after start codon. The TALEN-2 recognition sequence (purple line) begins after the 18 nucleotide spacer region. B: Chromatograms of sequence read from wild type (WT) and homozygous gjd2b^ncb215^ fish generated in this study indicate an insertion of nucleotide G (red arrowhead) within the Xho1 restriction site (blue stars). C: Representative gel image after Xho1 restriction digestion analysis of an 800bp amplicon (includes ∼300 bp upstream and 500 bp downstream sequences from the point of insertion) shows undigested and partially digested bands in the homozygous and heterozygous mutants respectively, compared to complete digestion in wild type siblings. D: Predicted amino acid sequences of various gjd2b mutant alleles generated in this study aligned with wild type Gjd2b sequence. gjd2b^ncb215^ is predicted to code for the first 6 amino acid residues of Gjd2b followed by nonsense sequence upto the 54^th^ amino acid position. Presence of a premature stop codon terminates translation at this position. E: Gjd2b-like immunoreactivity is reduced in PNs of gjd2b^ncb215^ compared to wild type larvae (Wilcoxon rank sum test; P=5.1283e-05). Number of images analyzed is indicated in parantheses. Staining was not completely abolished as the antibody also recognizes Gjd2a.

We next recorded mEPSCs from PNs in gjd2b^-/-^ larvae to determine if loss of Gjd2b leads to reduced glutamatergic synaptic contacts, as observed in the morphants (Figure 1). gjd2b^-/-^ larvae showed significant increase in mEPSC inter event intervals (Figure 4A, C), recapitulating the results observed after knock down of Gjd2b with SBMO (Figure 1C). Further, homozygous mutants also showed an increase in peak amplitudes (Figure 4A, B, D), and faster kinetics as revealed by decreases in rise times (Figure 4B and E) and decay time constants (Figure 4B and F). These results are consistent with the results obtained after knocking down Gjd2b with SBMO and point to a decrease in the number of glutamatergic synaptic contacts impinging on PNs.

**Figure 4:**
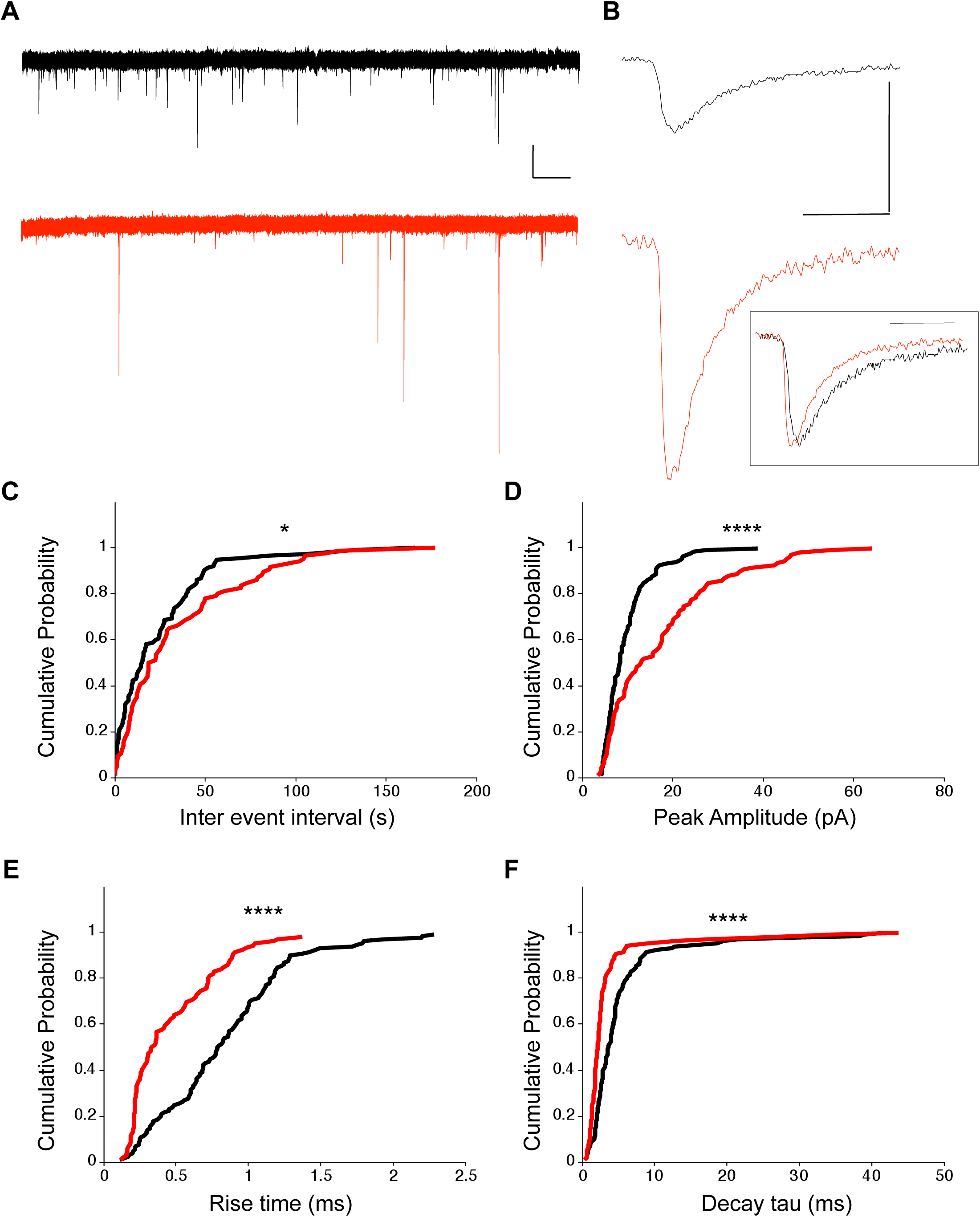
Knocking out Gjd2b also results in decrease of glutamatergic synaptic number. A: Representative mEPSC recordings from PNs of wild type (black trace) and gjd2b^-/-^ (red) larvae. Scale bars: x: 10ms; y:5pA. B: Average mEPSC shown on expanded time base recorded from wild type (black) and gjd2b^-/-^ (red) larvae. Scale bars: x: 5ms; y: 10pA. Inset: Scaled mEPSC to show faster rise time and decay time of mEPSCs in mutants. Scale bar: 5ms. C-F: Cumulative probability histograms reveal increased inter event intervals (C), increased peak amplitudes (D), decreased 10-90% rise times (E) and decreased decay time constants (F) of mEPSCs in gjd2b^-/-^ larvae (red lines) compared to wild type (black lines). N = 8 cells in wild type and 10 cells in gjd2b^-/-^ larvae. *P<0.05; ****P<0.0001; Mann-Whitney U test.

To finally confirm if the loss of Gjd2b indeed results in a decrease in synapse density, we quantified synapse density at the ultrastructural level. PNs send elaborate dendritic arbors into the molecular layer of the cerebellum where they make excitatory synapses with parallel fibers and climbing fibers and inhibitory synapses with axons of molecular layer interneurons. Transmission electron micrographs (TEM) were obtained from the molecular layer of the corpus cerebelli (CCe) of 7 dpf wild type and mutant larvae from 60 nm thick sections of the brain. Sections were taken at an interval of 1.2 µm to avoid oversampling the same synapses. Synapses were counted as membrane appositions with presynaptic vesicles on one side and electron-dense post-synaptic density on the other (Figure 5A). We found that the density of synapses per cubic micrometer was significantly lower in gjd2b^-/-^ larvae compared to wild type larvae (Figure 5B). To understand if Gjd2b also regulates the maturation of synapses, we calculated the synapse maturation index [20, 32] for individual synaptic profiles, measured as the ratio of the area occupied by vesicles to the entire area of the presynaptic terminal (Figure 5C). The distribution of synapse maturation indices was not different between wild type and mutant larvae (Figure 5D). In sum, these results indicate that Gjd2b is essential for the formation of synapses onto PNs, but once formed, the maturation of synapses is independent of Gjd2b.

**Figure 5:**
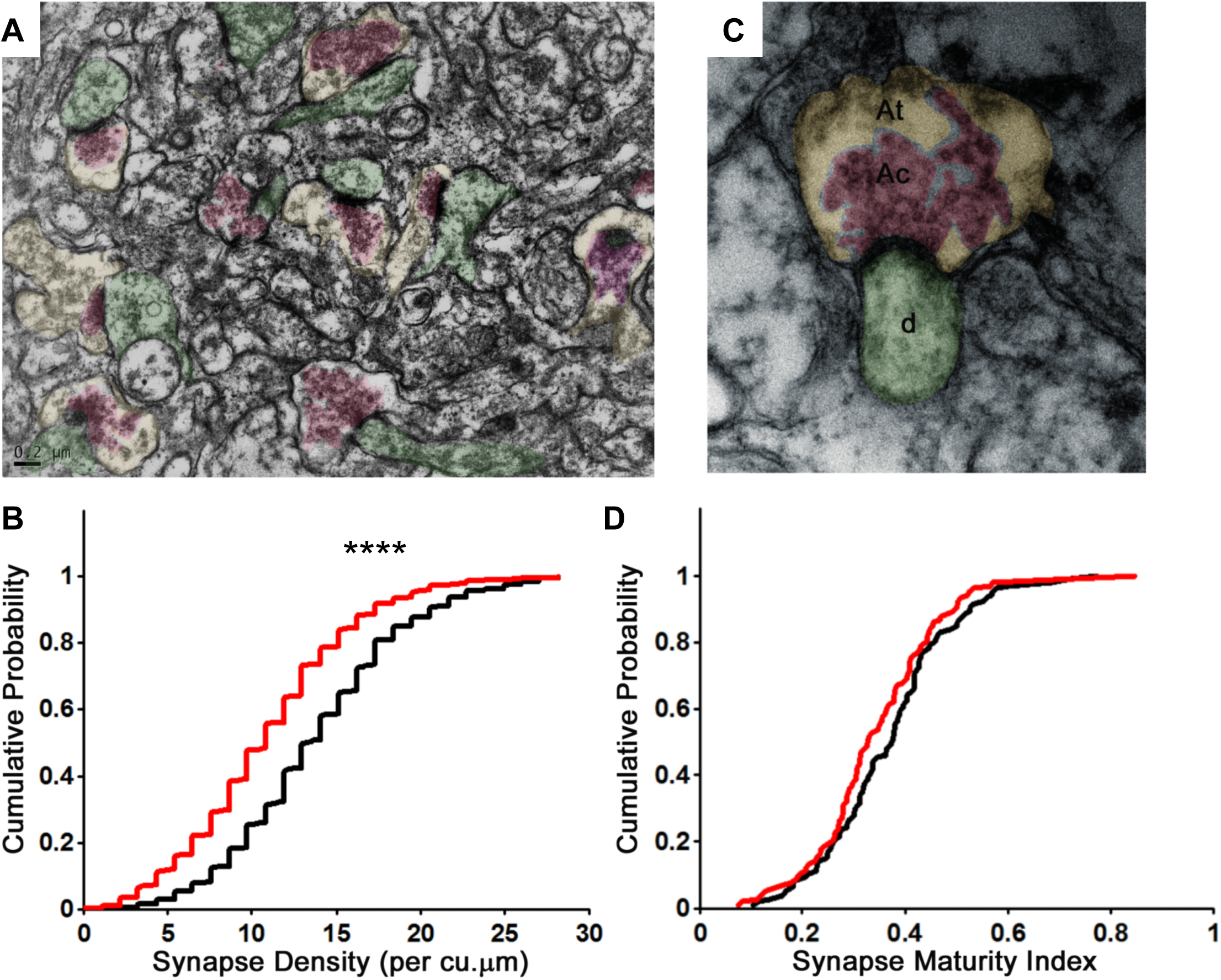
Knocking out Gjd2b leads to reduction in synaptic density in the cerebellar molecular layer. A: Transmission electron micrograph illustrating synapses identified using clustered vesicles (pink areas) in presynaptic terminals (yellow areas) and apposed post-synaptic density in dendritic profiles (green areas). B: Cumulative probability plot showing distribution of synapse density per cubic micrometer in wildtype sibling (black) and gjd2b^-/-^ larvae (red). 637 micrographs from 3 wild type larvae and 550 micrographs from 3 mutant larvae were analysed. ****P<0.0001; Mann-Whitney U test. C: Transmission electron micrograph at high magnification used for quantification of synapse maturity index. The area occupied by clustered vesicles (pink, Ac) was divided by the total area of the presynaptic terminal (yellow, At) to obtain the maturity index. D: Cumulative probability plot showing the distribution of synapse maturity indices in wild type sibling (black) and gjd2b^-/-^ larvae. 106 micrographs from 3 larvae each in wild type and mutant groups were analyzed. P = 0.12, Mann-Whitney U test.

The increase in peak amplitude combined with faster kinetics of mEPSCs in gjd2b^-/-^ mutants suggested that synapses in mutant PNs are placed electrotonically closer to the soma than in wild type larvae. This in turn implies that loss of Gjd2b could result in stunted dendritic arbors thereby resulting in the loss of distally located synapses. To understand if this is indeed the case, we labeled PNs of wild type and mutant larvae in a mosaic fashion and imaged them daily from 5 dpf till 8 dpf (Figure 6A, B), a period when PNs in larval zebrafish are growing and making synaptic connections [23, 24].

**Figure 6:**
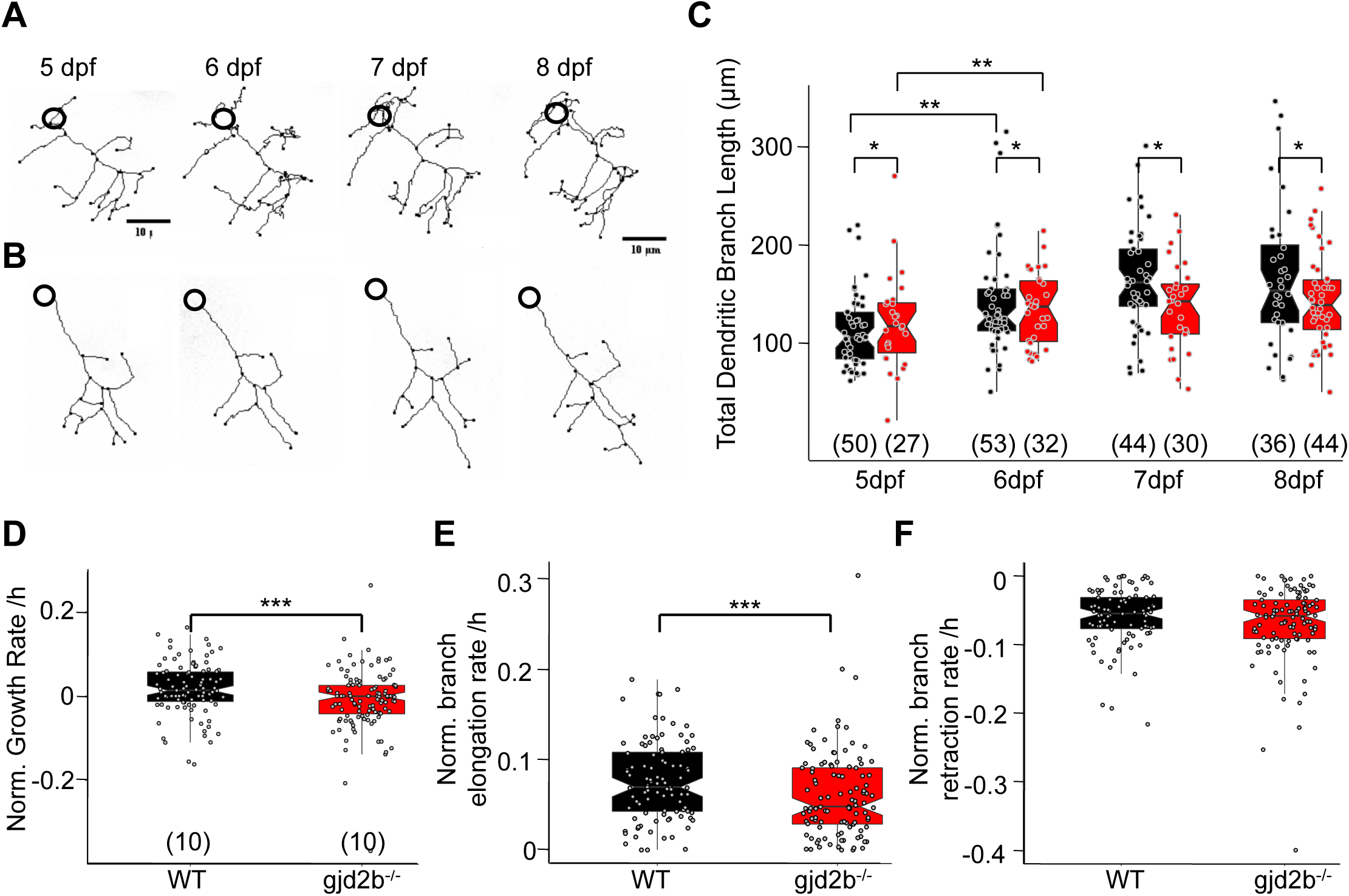
Dendritic arbor growth of gjd2b^-/-^ PNs is impaired. A. Wild type PN traces from 5-8 dpf. B. gjd2b^-/-^ mutant neuron traces from 5-8 dpf (Scale bar = 10 µm). C. Total dendritic branch length (TDBL) of WT (black) and gjd2b^-/-^ (red) PNs from 5-8 dpf. WT and gjd2b^-/-^ neurons show significant growth from 5-6dpf (p=0.0023). WT neurons show significantly higher TDBL values at all days (p=0.048). Statistical comparison was done using a general linear model with an inverse gaussian error distribution. Post-hoc comparisons were done with the emmeans package in R. D. Average rate of normalized hourly branch growth in wildtype and gjd2b^-/-^ PNs at 6dpf. Mutant neurons show a significantly reduced rate over 10 hours of observation (p=0.005). E. Rate of branch elongation in wildtype and mutant PNs at 6dpf. Mutant PNs show a significantly reduced rate of branch length elongations (p=0.022) F. Rate of branch length retractions in wildtype and mutant PNs at 6dpf. No significant difference is observed between the two groups (p=0.185). Statistical comparison for D-F was done using general linear models with gaussian error distribution. N’s (number of neurons sampled) are indicated in parantheses in C-E. See also Figure S3.

We measured the total dendritic branch length (TDBL) of PNs in wild type and mutant larvae at 5, 6, 7 and 8 dpf. Wild type PNs showed a significant increase in their TDBL from 5 dpf to 6 dpf, but not at later stages (Figure 6C). gjd2b^-/-^ PNs also showed a similar pattern of branch length growth, with an increase in TDBL from 5 to 6 dpf, followed by no significant increase until 8 dpf (Figure 6C). Post-hoc analysis revealed that there was a significant difference in TDBL between WT and gjd2b^-/-^ on all 4 days, with the mutant group having consistently lower values (Figure 6A-C). PN somata in wild type and gjd2b-/- larvae were comparable in diameter (Figure S3A). These analyses reveal that loss of Gjd2b leads to a stunted dendritic arbor resulting in fewer synapses placed proximal to the soma. To understand the dynamics that result in stunted and branched arbors, we imaged PNs every hour at 6 dpf for 10 hours. Mutant neurons had a significantly reduced growth rate compared to wild type PNs (Figure 6D). When dissected further into branch elongations and retractions, mutant neurons had a significantly lower rate of branch elongations than wild type neurons (Figure 6E), but their rates of branch length retractions were similar (Figure 6F). This suggests that Gjd2b regulates dendritic growth by promoting branch elongations and is unlikely to be involved in regulating branch retractions.

We next wished to determine which neurons are electrically coupled to PNs. Electroporation of single PNs with a combination of a high molecular weight dye (tetra-methyl rhodamine dextran) and a low molecular weight tracer (neurobiotin) failed to reveal any dye-coupled cells. However, when non-PNs were electroporated with neurobiotin, one or two PNs along with several non-PNs were detected (Figure S4, Table S1), indicating that PNs are likely to be coupled to other cerebellar cell types via rectifying junctions. To determine if the observed dendritic growth deficits in gjd2b^-/-^ mutant PNs are due to lack of Gjd2b in PNs specifically, we introduced Gjd2b tagged to mCherry into single PNs in gjd2b^-/-^ fish (Figure 7A). TDBL in mutant PNs expressing Gjd2b were significantly larger than mutant PNs and were rescued to wild type levels (Figure 7B). This suggests that the presence of Gjd2b in single PNs in otherwise Gjd2b null larvae is sufficient to guide dendritic arbor elongation. Heterotypic gap junctional channels formed by Gjd2b on PN membranes and a different connexin isoform in the coupled cell could mediate this process.

**Figure 7:**
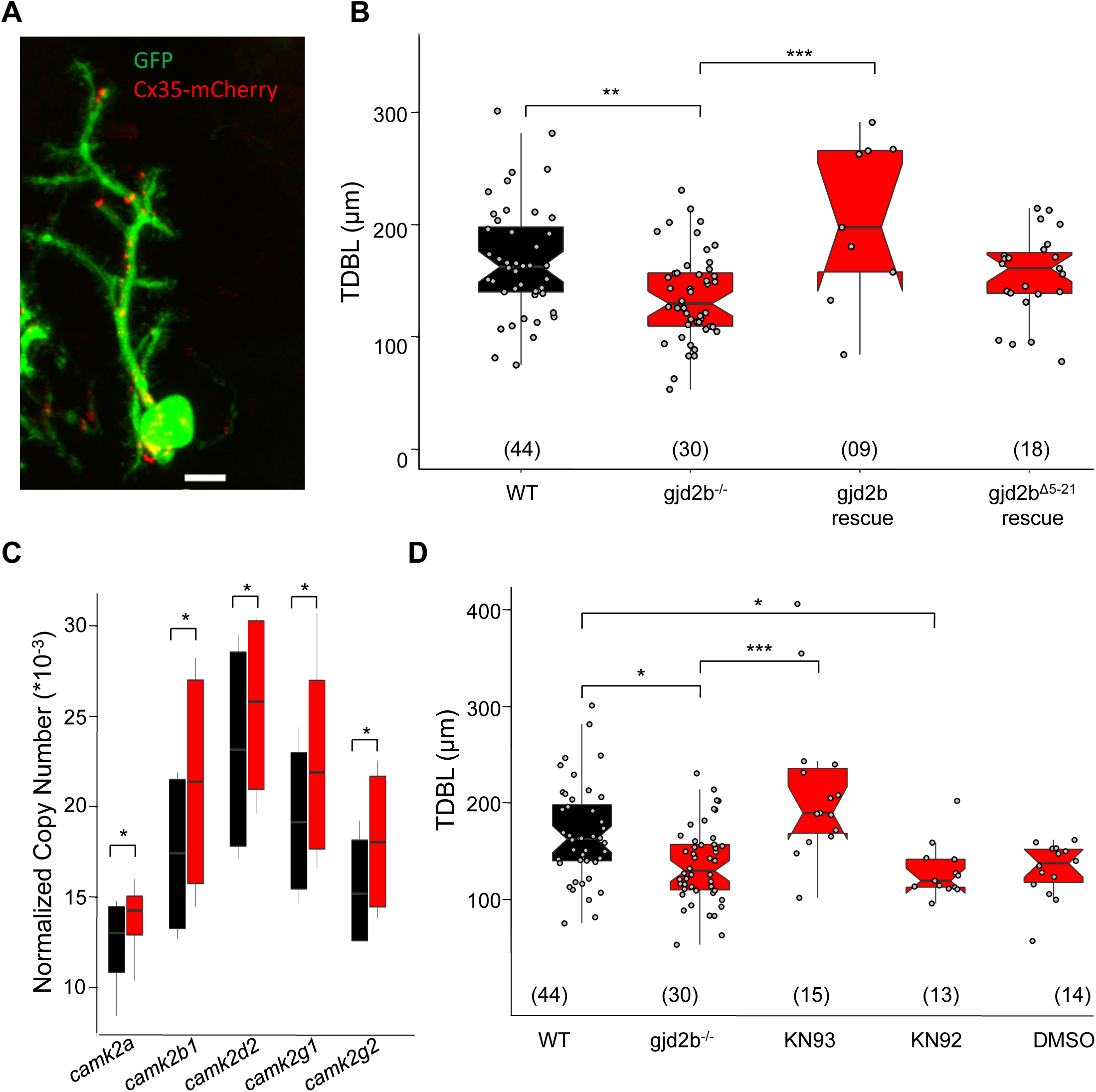
Expressing Gjd2b in PNs alone or inhibiting CaMKII is sufficient to rescue dendritic growth deficits. A. Representative image of a zebrafish PN expressing cytoplasmic GFP (green) and Gjd2b tagged with mCherry (red) at 7dpf (scale bar = 5 µm). B. Total dendritic branch length (TDBL) of wild type (WT), gjd2b^-/-^, gjd2b-rescue and gjd2b^Δ5-21^ rescue (pore dead variant) PNs at 7dpf. TDBL of gjd2b-rescue neurons showed no significant difference from the wild type group, but is significantly increased from that of mutant PNs. TDBL of Gjd2b^Δ5-21^ rescue PNs is not significantly different from that of gjd2b^-/-^ PNs. C. Copy number of CaMKII mRNA in WT and gjd2b^-/-^ larvae. The normalised copy number of CaMKII in the mutant group is significantly higher than the WT group (p<0.05) D. TDBL of WT, untreated gjd2b^-/-^, 1 µM KN-93 treated, 1 µM KN-92 treated and 1% DMSO treated gjd2b^-/-^ PNs at 7dpf. TDBL of only KN-93 treated gjd2b^-/-^ PNs are rescued to wild type levels (wt vs. KN-93, p = 0.23), whereas all other groups are significantly different and lower than wild type (Kruskal Wallis, post-hoc comparison with Mann Whitney test). N’s (number of neurons sampled) are indicated in parantheses in B and D. See also Figure S4 and Table S1.

To test if functional Gjd2b-mediated gap junctions are required for dendritic elaboration, we generated a construct coding for an N-terminal deleted version of Gjd2b (Gjd2b^Δ5-21^), as the N-terminus of connexins has been shown to be required for channel function but not for assembly into gap junctional plaques [33]. Expression of Gjd2b^Δ5-21^ in PNs of gjd2b larvae resulted in dendritic arbors that remained stunted and not significantly larger than mutant PNs lacking Gjd2b (Figure 7B). These results point to the need for functional gap junctional channels in regulating PN dendritic elaboration.

To further understand how Gjd2b-mediated gap junctions regulate synaptogenesis and dendritic growth, we focused on cytoplasmic binding partners of Gjd2b that are also known to play a significant role in these processes. The calcium and calmodulin dependent kinase II (CaMKII) has been shown to associate with Cx35/36-mediated gap junctions and modulate their function [34–37]. CaMKII is localized in dendrites and in spines and is a critical regulator of dendritic development [38–40]. We first asked whether expression levels of the various isoforms of CaMKII are altered in gjd2b^-/-^ larvae compared to wild type. To our surprise, we observed increased copy numbers for all isoforms of *camk2* we tested in gjd2b^-/-^ compared to wild type (Figure 7C). CaMKII has previously been shown to reduce dendritic dynamics and stabilize branches [38, 39]. To test if the stunted arbors observed in gjd2b^-/-^ PNs were due to premature stabilization of dendritic arbors, mediated by increased CaMKII activity, we inhibited CaMKII in gjd2b^-/-^ larvae using the drug KN-93 in the embryo medium. Incubation inKN-93 was able to rescue dendritic arbor lengths of PNs in gjd2b^-/-^ larvae to wild type levels (Figure 7D). No changes in dendritic arbor lengths were observed in larvae incubated in DMSO (vehicle control) or in KN-92, an inactive analog of KN-93 (Figure 7D). Taken together, these results show that conduction via Gjd2b-mediated electrical synapses is critical for proper glutamatergic synaptogenesis in PNs and that this process affects dendritic arbor growth by reducing the levels of CaMKII.

## Discussion

### Gap junctions in PNs

Gap junction proteins are widely expressed in the cerebellum of developing and adult vertebrates. Cell type specific transcriptome analysis in larval zebrafish revealed gjd2b expression in Purkinje neurons, granule cells, eurydendroid cells and Bergmann glial cells [41]. We found Gjd2b puncta localized to PN cell membrane and dye-coupling of PNs to other non-PN cells. At this time, the identity of the coupled cells is not known. In rodents, few studies have reported dye-coupling of PNs to other PNs or to other cell types in the cerebellar cortex [42–45]. Cx36, the mammalian homolog of Gjd2b has been shown to be expressed in the cerebellar cortex in the molecular layer and granule cell layer [46–48], although the mRNA was not found in Purkinje neurons [46]. From these studies, it appears that PNs in fish and mammals are electrically coupled to other neurons, while there may be species-specific differences in the molecular components of these electrical synapses. We suggest that despite differences in the molecular components, gap junctions on PNs could serve important developmental functions in all vertebrates.

### Regulation of chemical synapse formation by electrical synapses

Our results indicate that Gjd2b-mediated functional electrical synapses are important regulators of glutamatergic synapse formation and dendritic elaboration. These results are in agreement with earlier studies on innexin-mediated gap junctions in invertebrates. Knock-down of the innexin *inx1* in leech resulted in loss of electrical coupling between identified neurons at embryonic stages and decreased chemical synaptic strength between the same neurons at much later stages [18]. In neocortex of mice, sister neurons born from the same radial glia, make transient electrical synapses, which are required for the formation of excitatory connections between them [49, 50]. Mice lacking Cx36 exhibit reduced synaptic connectivity between mitral cells in the olfactory bulb [17]. Interestingly, though Cx36 is the predominant neural connexin, deficits in glutamatergic synapse formation have hitherto not been reported from other regions of the CNS, to the best of our knowledge. However, an increase in the number of inhibitory synapses was observed in Cx36^-/-^ mice in thalamocortical relay neurons with a concomitant decrease in their dendritic complexity [51]. Using morpholino-mediated knock down and knock out approaches, we show that glutamatergic synapse number is decreased significantly when Gjd2b/Cx35b is perturbed. The decrease in synapse number was seen using both structural (transmission electron microscopy) and functional (electrophysiology) assays. These results provide direct evidence for the role of electrical synapses in instructing glutamatergic synapse formation in the central nervous system.

### Dendritic development of PNs

PNs exhibit one of the most elaborate and beautiful dendritic arbors known and a number of studies have examined factors that determine arbor structure in PNs. In rodents, PN dendritic development occurs over a period of one month after birth and involves multiple steps. *In vivo*, rodent PNs undergo morphological changes soon after they reach the Purkinje cell layer. They retract their simple fusiform dendrites and then acquire a stellate morphology with multiple dendrites sprouting from their somata [52–54]. This has also been observed in zebrafish PNs ([55] and Sitaraman and Thirumalai, unpublished observations). Later, one of these becomes the primary dendrite and the others are retracted. This process involves the localization of Golgi organelles at the base of the primary dendrite and is mediated by an atypical PKC [55]. These early steps occur roughly before 4 dpf in zebrafish and before P10 in rodents. The early remodeling is mainly dependent on intrinsic factors and the overall architecture of PN dendrites is maintained even in the absence of afferent input or activity. In the second post-natal week, the apical dendrite of rodent PNs is spiny and undergoes growth and branching to occupy the molecular layer in a planar manner. By P20 in mice and P30 in rats, PN dendrites have achieved their maximal length [52, 53]. From our results, it appears that significant growth of PN dendritic arbors occurs in zebrafish between 5 and 8 dpf. At these stages, the neurons are spiny and receive parallel and climbing fiber inputs [25]. We also observed that at these stages, the dendritic branches are dynamic and undergo elongations and retractions. In rodents, lack of afferent input during this phase results in abnormal orientation, reduced size and lack of higher order branches in PN dendritic arbors [56, 57]. In both slice cultures and dissociated cell cultures, blockade of glutamatergic transmission reduces dendritic arbor size of PNs [58, 59]. We observed that dendritic arbors of gjd2b^-/-^ PNs were smaller compared to wild type even at 5 dpf and stayed smaller at least until 8 dpf. During these stages, they also received fewer glutamatergic mEPSCs. We suggest a ping-pong mechanism by which the smaller dendritic arbor restricts the number of functional synapses that can be formed and the reduced afferent input in turn restricts further dendritic growth. It is also likely that signaling via gap junctions promotes dendritic arbor growth directly, independent of their actions on chemical synaptogenesis.

### Role of CaMKII in gap-junction mediated PN development

We could rescue dendritic arbor growth deficits in Gjd2b mutant zebrafish by expressing full length Gjd2b in single PNs. In addition, we found that expressing an N-terminal deleted, pore-dead version of Gjd2b could not rescue the dendritic growth deficit. These results suggest that conduction of signaling molecules through Gjd2b-mediated gap junctions regulates dendritic arborization. Further, Gjd2b mutants show increased expression levels of α, β1, δ2, γ1, and γ2 isoforms of CaMKII and inhibition of CaMKII restores the dendritic arbor lengths in mutants to wild type levels. In sum, these results lead us to propose that passage of signaling molecules into PNs via Gjd2b-mediated gap junctions keeps CaMKII levels low enough to allow dendritic branches to grow and elongate. When CaMKII levels and/or activity increases, dendrites are stabilized at their mature lengths. Lack of gap junctional signaling leads to premature stabilization of dendritic branches leading to stunted growth. In *Xenopus* tectum, immature neurons with simple arbors and low levels of CaMKII, continue to grow while mature neurons have high levels of CaMKII and their dendritic structure is more or less stable. Expression of constitutively active CaMKII in tectal neurons causes them to grow slower and have relatively less dynamic arbors. In addition, inhibition of CaMKII in mature neurons causes them to grow at a higher rate [38]. More recently, a human CAMK2A mutation, isolated from an ASD proband, was shown to cause increased dendrite arborization, when the mutant CAMK2A was introduced into cultured mouse hippocampal neurons [60]. Our results are consistent with these earlier findings and point to a stabilizing role for CaMKII in PN dendritic arbor elaboration. The mechanisms by which signaling via Gjd2b gap junctions regulates CaMKII levels will have to be investigated in future experiments.

## Materials and Methods

### Zebrafish and animal husbandry

All experiments were performed using Indian wild type zebrafish. Institutional Animal Ethics and Bio-safety committee approvals were obtained for all procedures adopted in this study. Larvae and adults were reared using standard procedures[61].

### Generation of transient transgenic larvae

To label single PNs for some of the experiments, single-celled embryos were microinjected with one of the following constructs along with Tol2 transposase mRNA[62]: aldoca:gap43-Venus[55] (gift from Prof. Masahiko Hibi, Nagoya University, Japan); Ca8-cfos: RFP[63] (gift from Dr. Hideaki Matsui, Niigata University, Japan); Ca8-cfos:GFP; Ca8-cfos:Gjd2b-mCherry; Ca8-cfos:Gjd2b^Δ5-21^- mCherry. Ca8-cfos: GFP was constructed by amplifying Ca8-cfos from the parent plasmid and ligating it with sequences coding for GFP. To construct the last two plasmids, full length Gjd2b and Gjd2b^Δ5-21^ were first amplified from total cDNA using the following primers: Gjd2b forward: ^5’^GATCGGTACCTCCGAATGAACAGCCAT^3’^; Gjd2b reverse: ^5’^TAGCGCTAGCAACGTAGGCAGAGTCACTGG^3’^; Gjd2b^Δ5-21^ forward: ^5’^ATTGCCATGGGGGAATGGATTGGGAGGATCCTGCTAAC^3’^; Gjd2b^Δ5-21^ reverse: ^5’^TAGCGCTAGCAACGTAGGCAGAGTCACTGG^3’^. The amplified regions were restriction digested using *NcoI* and *NheI* and ligated with Ca8-cfos on the 5’ end and mCherry at the 3’ end to generate the respective plasmids. Microinjected embryos were reared in embryo medium containing 0.003% of 1-phenyl-2-thiourea (PTU) for imaging experiments.

### Morpholino antisense oligonucleotide mediated knockdown of Cx35

Morpholino antisense oligonucleotides (Gene Tools LLC) were designed to bind at the junction between exon1 and intron1 of the gjd2b mRNA to block proper splicing of the mRNA (SBMO; Figure S2A). Control morpholinos (CTRL) were designed to incorporate mismatches at 5 positions within the gjd2b recognition sequence. The morpholino sequences were:

SBMO: 5’ ACAACACTTTTTCCCCTCACCTCCC 3’

CTRL: 5’ ACTAGACTTATTCCCGTGACCTCCC 3’

Either SBMO or CTRL were injected into single-celled zebrafish embryos at 0.05 pmoles per embryo.

### Generation of gjd2b^-/-^ zebrafish

Transcription activator like effector nucleases (TALENs) recognizing nucleotide sequences near the start codon of gjd2b gene were used to generate the gjd2b mutant (*gjd2b^-/-^*) lines of zebrafish used in this study (Figure S3). A pair of TALEN vector constructs were designed and assembled to generate gjd2b-TALEN-1 and gjd2b-TALEN-2, which bind the plus and minus strands of *gjd2b* gene (Figure S3A), respectively, by following published protocols[64]. These TALEN sequences were later moved to a pTNT^TM^ (Promega Corp, Madison, WI) vector. TALEN mRNAs that encode gjd2b-TALEN-1 and gjd2b-TALEN-2 proteins, were synthesized *in vitro* from the above vectors using the T7-mMessage mMachine kit (Life Technologies, Thermo Fisher Scientific, USA) and micro-injected into one cell stage wildtype zebrafish embryos at a concentration of 50ng/µl of each mRNA. TALENs were designed such that the spacer region incorporated an *Xho1* restriction site enabling an easy screen for mutations in this locus. Upto ten embryos were taken from every clutch of TALEN-injected embryos and screened for the presence of mutations using *Xho1* restriction digestion. Clutches of embryos that showed a high percentage of mutants were then grown up into adults. The TALEN-injected founder generation (F0) was raised in the facility and the adult fish were out-crossed with wild type fish to get heterozygous F1 progeny. These were screened for germline transmission of mutations in the *gjd2b* locus. Whole embryos or adult fish tail clips were screened using *Xho1* restriction analysis of 800 bp DNA band around the TALEN-target site followed by sequencing of this region. F1 heterozygous mutants (*gjd2b^+/-^*) were inbred to obtain F2 wild type siblings, heterozygotes and homozygotes (*gjd2b^-/-^*).

### Whole-mount immunohistochemistry

Whole mount immunofluorescence using anti-Cx35/36 antibody was performed as described in [16]. Briefly, 5-7 dpf larvae were anaesthetized in 0.01% chilled tricaine (Sigma-Aldrich) and then fixed overnight at 4°C in 4% paraformaldehyde (Alfa Aesar, Thermo Fisher Scientific, UK), followed by several washes in 0.1M Phosphate buffered saline (PBS) at room temperature. The eyes, jaws and yolk sac were carefully dissected out and the skin covering the brain was peeled to expose the brain. Dissected larvae were kept overnight in 5% normal donkey serum and 0.5% Triton-X100 in 0.1M PBS (PBST) and at 4°C. Then they were incubated in mouse anti-Cx35/36 antibody (MAB3045, EMD Millipore, Merck, USA) at a dilution of 1:250 in blocking solution for 48 hours. After several washes in PBST, larvae were incubated overnight in anti-mouse Alexa Fluor 488 antibody (Invitrogen) at a dilution of 1:500 in blocking solution. Following this, the larvae were washed in 0.1M PBS several times and then mounted between two coverslips using Prolong Gold anti-fade reagent (Molecular probes, Life technologies, Thermo Fisher Scientific, USA) and stored in the dark at 4°C until imaging.

### Confocal imaging and image analysis

Images were acquired on Zeiss LSM 780 or Olympus FV1000 confocal laser scanning microscopes using a 63X oil-immersion objective. Imaging parameters and settings were the same for all larvae imaged. Dorsal most regions of the cerebellum were imaged. Two images were taken from each animal, one in each hemisphere. Image analysis was done using Fiji (https://fiji.sc/)[65]. After background subtraction (rolling ball method), median filter was applied to remove salt and pepper noise from images. Average intensity of the z-projections of image stacks was then measured and normalized with respect to the intensity of uninjected (Figure S2) or wild type animals (Figure S3). These were then plotted using R statistical software (https://www.r-project.org).

### Transmission electron microscopy and image analysis

Zebrafish larvae at 7 dpf were fixed overnight in 4% paraformaldehyde and 2.5% glutaraldehyde prepared in EM buffer (70mM sodium cacodylate and 1mM CaCl_2_, pH 7.4). Secondary fixation was done with 1% OsO_4_ made in EM buffer for 90 minutes on ice. For additional contrast, samples were incubated in aqueous 2% Uranyl acetate for 1hr at room temperature. Samples were dehydrated serially in 30%, 50%, 70%, 90% and finally in absolute ethanol for 15 minutes each. After two changes of ethanol (15’ each), samples were incubated in acetone with two changes for 10 minutes each. Samples were infiltrated with Epon812-Araldite resin mix and acetone in the ratio of 1:3, 1:1, 3:1 and in pure Epon812-Araldite mix for 2hrs, overnight, 2hrs and overnight respectively. Samples were then embedded in the Epon812-Araldite resin mix for polymerization at 60°C for 48hrs. After polymerization, thick transverse sections were cut using a glass knife till the region of interest was obtained. Then, 60nm ultrathin sections were cut from the anterior end of the cerebellum on an ultramicrotome (Power Tome-PC, RMC Boeckeler) by using 3.5 size Ultra 45^0^ diamond knife (Electron Microscopy Sciences, USA). Sections were collected on Formavar/Carbon 2X1mm copper or Nickel slot grids (FCF 2010-Cu/Ni, Electron Microscopy Sciences, USA) for imaging. Images were acquired on FEI TECNAI T12 G^2^ Spirit BioTWIN transmission electron microscope. For counting synapses and for measuring synapse maturity indices, images were taken at 30K and 50K magnification respectively. Total volume of a single micrograph imaged was 0.924 cubic micrometers. Images were analysed on Fiji. Analysis was done blind to the genotype.

### Electrophysiology

Whole cell patch clamp recording from larval Purkinje neurons was performed as described in [25]. Briefly, 7 dpf larvae were anaesthetized in 0.01% MS222 and then pinned onto a piece of Sylgard (Dow Corning) glued to a recording chamber. Then larvae were submerged in external saline (134mM NaCl, 2.9mM KCl, 1.2mM MgCl_2_, 10mM HEPES, 10mM Glucose, 0.01 D-tubocurarine, 2.1mM CaCl_2_,; pH 7.8; 290mOsm;). The cerebellum was exposed by peeling the skin from the top of the head. Cells were viewed using 63X water immersion objective of a fixed stage compound microscope (Nikon Ni-E or Olympus BX61WI). Patch pipettes were pulled from borosilicate glass (OD: 1.5mm; ID: 0.86mm; Warner Instruments) on a Flaming-Brown P-97 pipette puller (Sutter Instruments), filled with pipette internal solution and had a resistance of 10 to 12 MΩ. For miniature excitatory post synaptic current recording, cesium gluconate pipette internal solution was used (115mM CsOH, 115mM Gluconic acid, 15mM CsCl, 2mM NaCl, 10mM HEPES, 10mM EGTA, 4mM Mg-ATP; pH 7.2 and osmolarity 290 mOsm). mEPSCs were recorded after bath application of TTX (1µM), Gabazine (10µM) and APV (40µM). Cells were held in voltage clamp at −65mV for all recordings. In a few cells, the holding potential was varied to measure reversal potential for the mEPSCs. Pipettes also contained sulforhodamine dye (Sigma) and only those cells filled with the dye at the end of the recording and showing Venus expression were considered for further analysis. In addition, cells whose series resistance varied by more than 20% during the recording session or those that had input resistances lower than 1 GΩ were excluded.

For evoked synaptic current recording, potassium gluconate based internal solution was used (115mM K gluconate, 15 mM KCl, 2 mM MgCl_2_, 10 mM HEPES, 10 mM EGTA, 4 mM Mg-ATP; pH: 7.2; 290 mOsm). Evoked synaptic currents were recorded by stimulating climbing fibres (CFs) with a bipolar electrode (FHC, Bowdoin, ME, USA). Stimulus strength was gradually increased until no failures occurred. Paired pulse ratio was measured by stimulating CFs at various inter stimulus intervals (ISIs) ranging from 35ms to 550ms and calculating the ratio of the second EPSC to the first.

Signals were acquired using Multiclamp 700b amplifier and digitized with Digidata 1440A digitizer (Molecular Devices). Data analysis was done offline in Clampfit 10.2 (Molecular Devices) and statistical analysis was performed using custom scripts written in Matlab (The Mathworks, Natick, MA) and in R. mEPSCs were analysed blind to the experimental group.

### In vivo time lapse imaging and analysis

For daily imaging, 5 dpf larvae with sparse fluorescent protein labeling of PNs were anaesthetized in 0.001% MS222 (Sigma) for 1 minute. They were then mounted in 1.5% low gelling agarose (Sigma) on a custom made confocal dish with their dorsal side towards the coverslip and covered completely with embryo medium. Purkinje neurons in embedded larvae were imaged using a Leica SP5 point scanning microscope. Neurons were chosen based on their morphology, level of RFP/GFP expression and traceability. Imaging parameters were kept constant across samples. Post imaging, the larvae were released from agarose and allowed to recover in embryo medium for 24 hours. They were imaged again at 6, 7 and 8 dpf. For the hourly imaging, Purkinje neurons in 6 dpf larvae were imaged at 1-hour intervals for 10 hours. For gjd2b and gjd2b^Δ5-21^rescue experiments, 7dpf larvae with both GFP and punctate mCherry expression in PNs were anaesthetised and embedded in LGA. Purkinje neurons in embedded larvae were imaged using an Olympus FV3000 confocal microscope. Neurons were traced using the Simple Neurite tracer plugin in Fiji and their total dendritic branch length (TDBL) and total dendritic branch number (TDBN) were calculated.

For analysing the daily imaging data, TDBL and TDBN of the two genotypes, across days, were statistically compared using Generalized linear models (Inverse Gaussian and Poisson, respectively). For post hoc analysis, the emmeans package in R was used to compare the TDBL and TDBN of consecutive days within each genotype and that of same days across the two genotypes. Hourly growth rates, elongation rates and retraction rates of wild type and mutant PNs were compared using generalized linear models with Gaussian error distribution. For analysing the effects of gjd2b and gjd2b^Δ5-21^ rescue, the TDBL and TDBN across groups were statistically compared using ANOVA and post hoc Tukey HSD. Finally, to test the effects of CaMKII inhibition, the TDBL across groups was statistically compared using Kruskal Wallis and post hoc Mann-Whitney test.

### Drug treatments

Larvae were treated with one of the following drugs from 4 dpf till 7dpf: 1 µM KN-93 (Sigma-Aldrich; Catalog number 422708); 1 µM KN-92 (Sigma-Aldrich; Catalog number 422709); 1% DMSO (Vehicle control). Stock solutions of drugs were made in DMSO and diluted to final concentration in embryo medium.

### Total RNA extraction and Gene expression analysis

Larval whole brains (n=10 in each group) were dissected in external saline on 6 dpf. Dissected brains were pooled in each replicate for RNA extraction from corresponding wild type and *gjd2b^-/-^* backgrounds. A total of four such biological replicates per background were obtained, where each replicate from wild type background was paired with a replicate group from *gjd2b^-/-^* background, which were born, raised and processed for RNA extraction at same respective time points. Total RNA was extracted from whole brain of 6 dpf larval zebrafish using RNeasy micro plus RNA extraction kit (Qiagen, Germany). Total RNA (2ng/µl) from such replicates was used for cDNA synthesis using Superscript III reverse transcription kit according to manufacturer’s protocol (Invitrogen, USA). Absolute copy numbers were obtained by droplet digital PCR (ddPCR) using Evagreen mastermix (Biorad, USA) and gene-specific primers that were previously described for zebrafish genes[66] (*camk2a, camk2b1, camk2b2, camk2d1, camk2d2, camk2g1, camk2g2*, and a reference control *ef1*α). Absolute quantification in the form of number of copies per μL for each *camk2* isoform was normalized with the control *ef1*α copy numbers in each replicate. These normalized values were then used to compare gene expression levels for each *camk2* isoform between wild type and *gjd2b^-/-^* backgrounds. Statistical analysis was done creating two linear mixed models: one, with an interaction term between the genotype and the isoforms, and second, without such an interaction term. These two models were not significantly different from each other. Therefore, the model without interaction was used for further analysis. The emmeans package in R was then used to compare the normalized copy numbers of the wild type and *gjd2b^-/-^* backgrounds across all the isoforms.

## Acknowledgements

The authors would like to thank the following sources of funding support: Wellcome Trust-DBT India Alliance Intermediate and Senior fellowships (VT), Department of Biotechnology (VT), Science and Engineering Research Board, Department of Science and Technology (VT), Department of Atomic Energy (VT), CSIR-UGC fellowship (SJ), NCBS-TIFR graduate student fellowship (SS, VA and SJ), SERB Post Doctoral Fellowship (GY). The authors would also like to thank Prof. Masahiko Hibi for the aldoca construct and Dr. Hideaki Matsui for the Ca8 enhancer construct. Further thanks are also due to Mr. Sriram Narayanan and Ms. Shivangi Verma for technical assistance, and Mr. T.P. Jagadeesh for the maintenance of our fish lines. In addition we would like to thank the NCBS-TIFR Genomics facility and the Central Imaging and Flow Facility for support.

