## Supplementary Figures for "Gjd2b-mediated gap junctions promote glutamatergic synapse formation and dendritic elaboration in Purkinje neurons"

Supplementary Figure S1

A

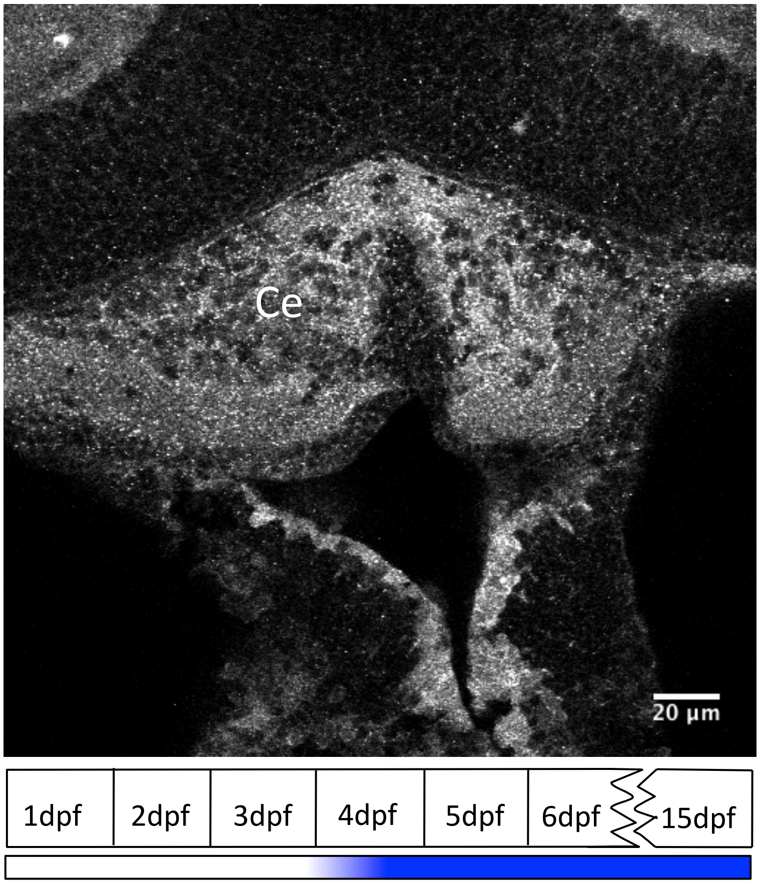


B

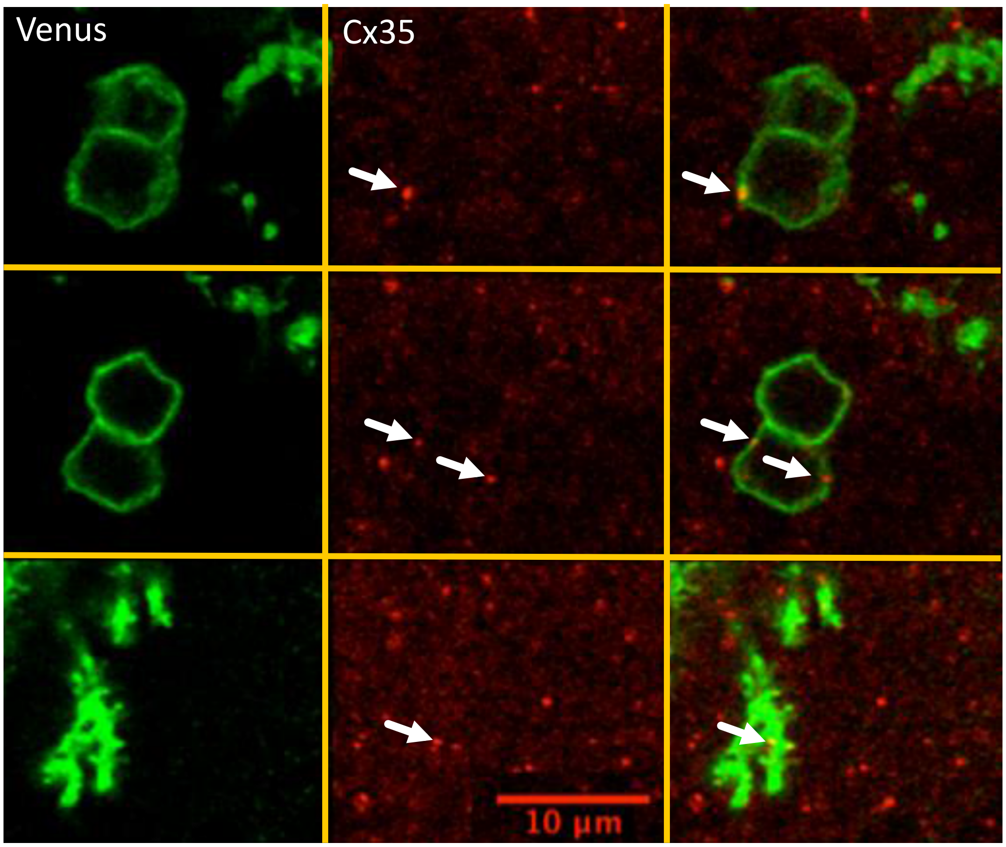

### Supplementary Figure S1

**Supplementary Figure S1: Connexin 35 (Cx35) is expressed in the cerebellum in PNs.** A: Cx35 immunoreactivity in the cerebellum (Ce) of 6dpf larvae. Schematic in the bottom shows the timeline for appearance of Cx35 immunoreactivity in the cerebellum of developing zebrafish. Adapted from Jabeen and Thirumalai, 2013. B: PNs labeled with aldoca: gap43-Venus (green) also show Cx35 immunoreactive puncta (red) in their somata (top two rows) and dendrites (bottom row). Related to Figures 1 and 2.

### Supplementary Figure S2

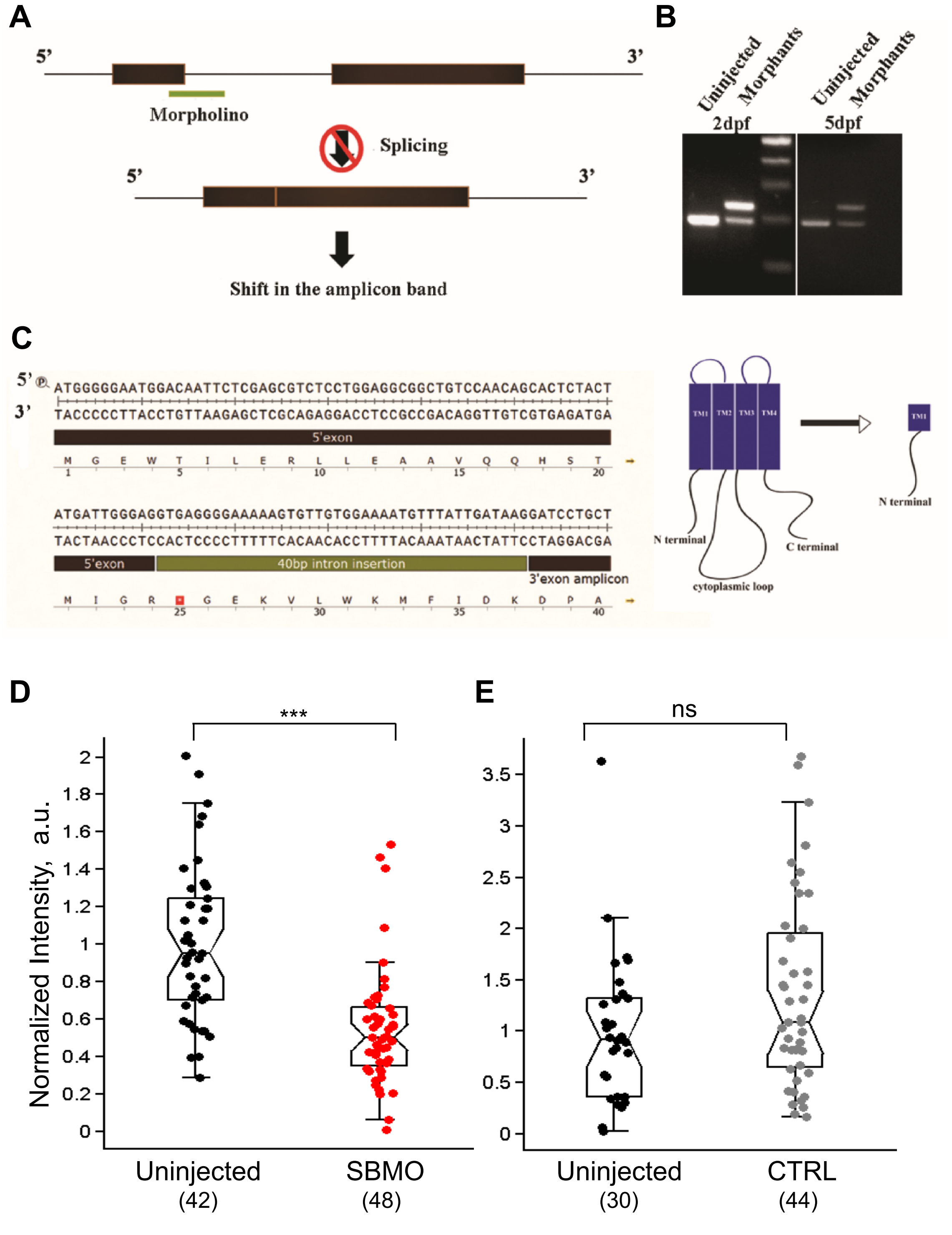


**Supplementary Figure S2: Effective knock-down of Gjd2b after injection of splice-blocking morpholino.** A: Schematic indicating the binding location of SBMO at the junction of exon1 and intron1 of gjd2b and blockade of normal splicing out of intron1. B: RT-PCR reveals a single amplicon in uninjected larvae but two amplicons, one of higher apparent molecular weight corresponding to the mis-spliced product, appearing in SBMO larvae at 2 dpf and 5dpf. C: Sequence of the mis-spliced gene product with a premature stop codon indicated by the red square. Predicted sequence of translated polypeptide is shown below and truncated Gjd2b with domain structure shown on the right. D: Reduced intensity of Gjd2b immunostaining in the cerebellum of SBMO larvae compared to uninjected larvae ($P<0.001$, Mann-Whitney test). E: Injection of a control morpholino with 5-base mismatch with target sequence (CTRL) does not alter Gjd2b immunostaining intensity in the cerebellum ($P= 0.14$, Mann-Whitney test). Number of images sampled in each group is indicated in parantheses at the bottom. Related to Figures 1 and 2.

Supplementary Figure S3

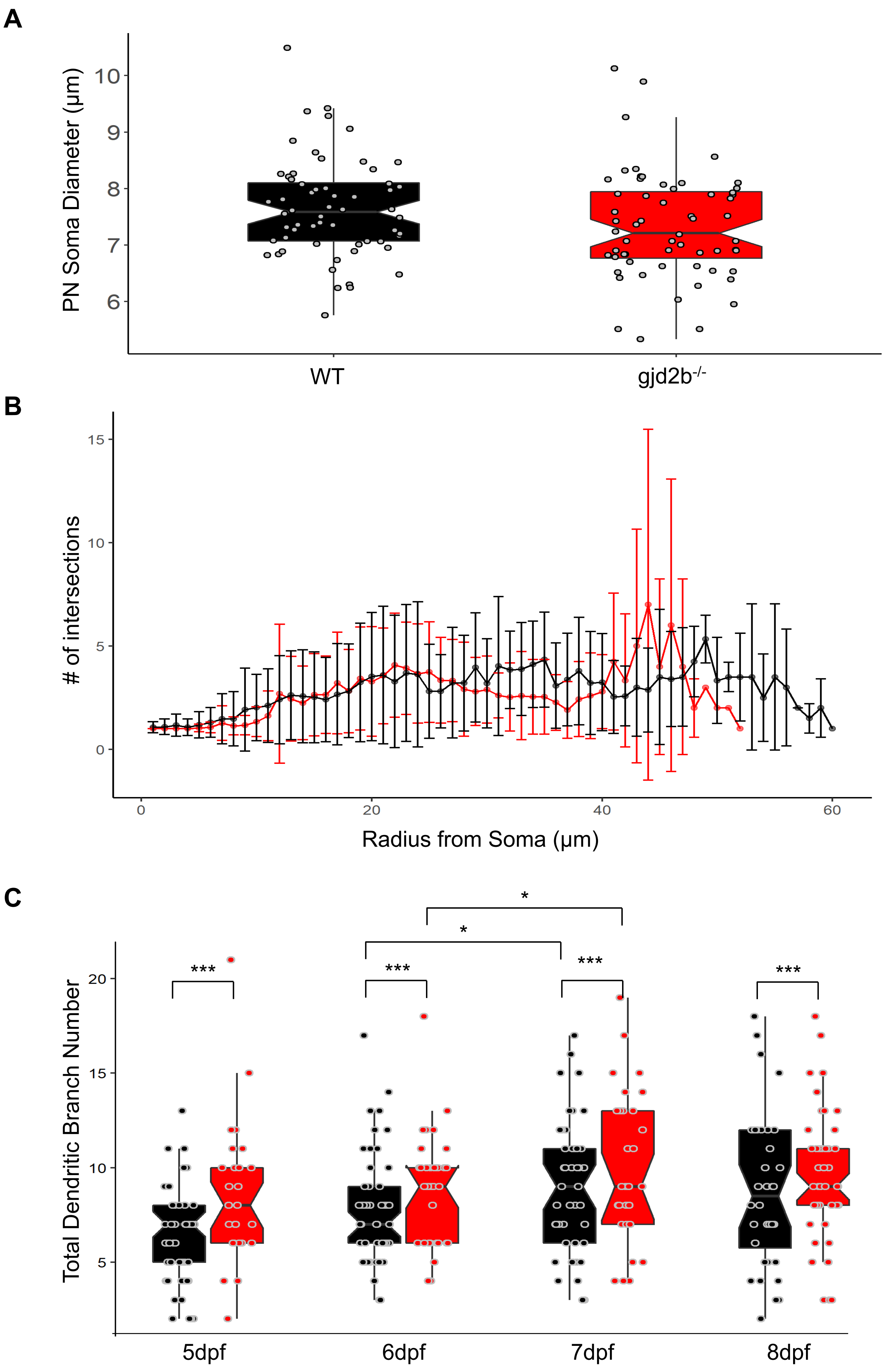

### Supplementary Figure S3

**Supplementary Figure S3: Additional morphometric comparison of PNs from wild type and gjd2b$^{-/-}$ larvae.** A. Total dendritic branch number (TDBN) for wildtype (black) and gjd2b$^{-/-}$ (red) PNs from 5-8dpf. WT and gjd2b$^{-/-}$ neurons show significant increase in branch number from 6-7dpf (p=0.0123). Gjd2b$^{-/-}$ neurons show significantly higher TDBN values at all days (p=0.0002). [Wildtype N(5dpf)=50 neurons; N(6dpf)=53 neurons; N(7dpf)=44 neurons; N(8dpf)=36 neurons] [gjd2b$^{-/-}$ N(5dpf)=27 neurons; N(6dpf)=32 neurons; N(7dpf)=30 neurons; N(8dpf)=44 neurons] B. Sholl analysis for wild type (Black; n=43) and gjd2b$^{-/-}$ (Red; n=29) neurons at 7dpf showing the average number of dendritic intersections at 1 μm radius steps. C. Soma diameter of wildtype (black) and gjd2b$^{-/-}$(red) PNs do not show any significant difference. [n(WT)=44, n(gjd2b$^{-/-}$)=30] (GLM with inverse gaussian error distribution, p=0.054). Related to Figure 6.

### Supplementary Figure S4

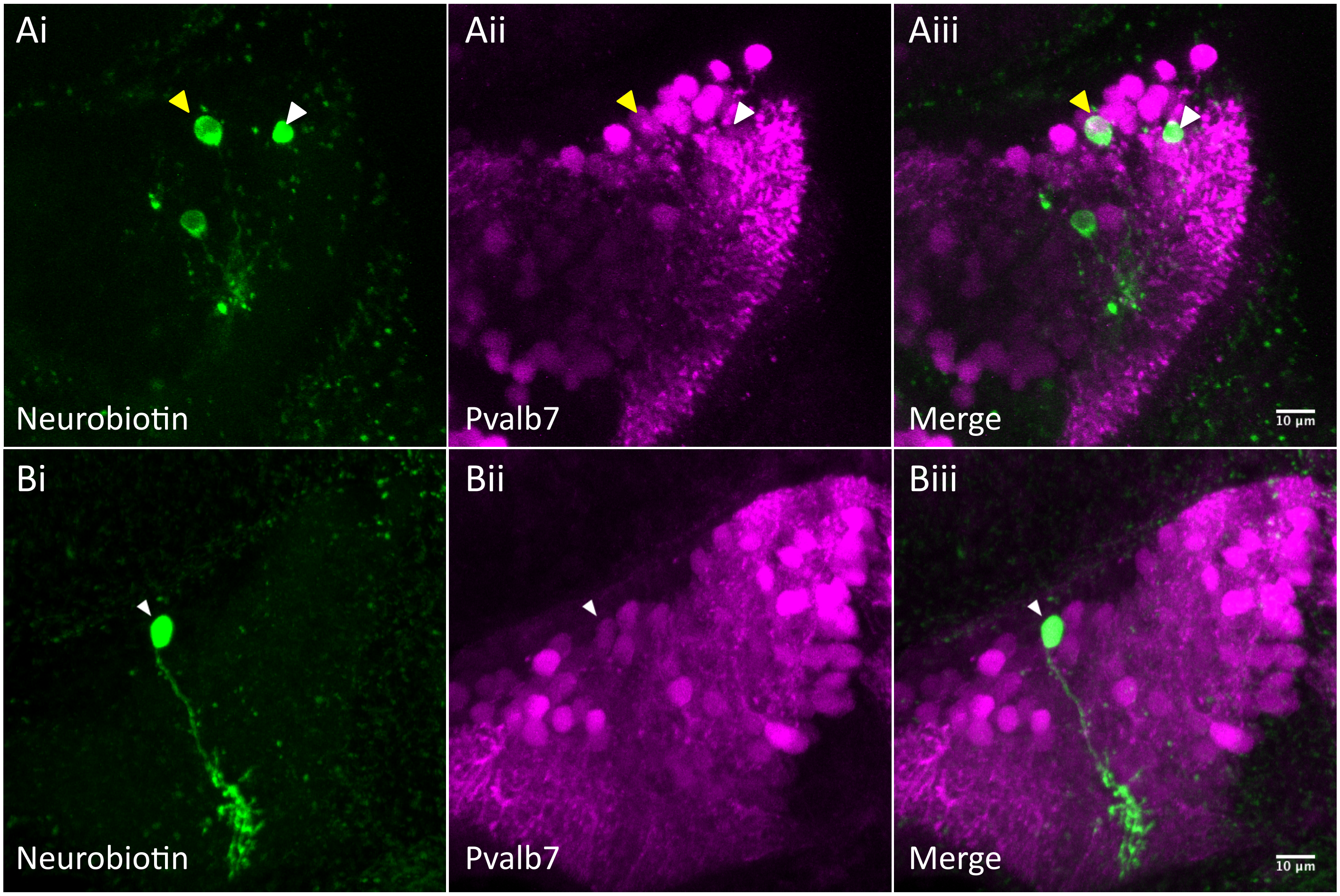


**Figure S4: PNs are dye coupled to other cerebellar cell types via putative rectifying junctions.** Ai-Aiii: Electroporation of neurobiotin (green) into a non-PN reveals several dye-coupled neurons of which one neuron is also Pvalb7 positive (magenta), indicating that it is a PN. Bi-Biii: Electroporation of neurobiotin into a PN reveals no other dye-coupled neurons.See also Table S1. Related to Figure 7. See also Table S1.
