## Supplementary Table S1 for "Gjd2b-mediated gap junctions promote glutamatergic synapse formation and dendritic elaboration in Purkinje neurons"

**Table S1: Single cell electroporation into PNs or non-PNs to identify electrically-coupled neurons.** When PNs were electroporated (animal nos. 1 through 8), no other dye-coupled neurons were detected. When non-PNs were electroporated (animal nos. 9 through 13), several other dye-coupled cells were seen, of which a couple were PNs. See also Figure S4.

| Electroporated Cell | | | No. of Dye Coupled Neurons | Dye coupled PNs |
| --- | --- | --- | --- | --- |
| Animal No. | PN | Non-PNs |  |  |
| 1 | 1 | 0 | 0 | 0 |
| 2 | 1 | 0 | 0 | 0 |
| 3 | 1 | 0 | 0 | 0 |
| 4 | 1 | 0 | 0 | 0 |
| 5 | 1 | 0 | 0 | 0 |
| 6 | 1 | 0 | 0 | 0 |
| 7 | 2 | 0 | 0 | 0 |
| 8 | 2 | 0 | 0 | 0 |
| 9 | 0 | 1 | 7 | 2 |
| 10 | 0 | 1 | 7 | 2 |
| 11 | 0 | 1 | 13 | 2 |
| 12 | 0 | 1 | 7 | 2 |
| 13 | 0 | 1 | 5 | 2 |
